# Intracortical Microstimulation Induces Rapid Microglia Process Convergence

**DOI:** 10.1101/2025.03.13.643075

**Authors:** Nathaniel P. Williams, Anna M. Kelly, X. Sally Zheng, Alberto L. Vazquez, X. Tracy Cui

## Abstract

Intracortical microstimulation (ICMS) has demonstrated the potential to restore vision and hearing by stimulating relevant cortical regions in both animals and humans, offering significant clinical promise for sensory restoration. While the neuronal response to ICMS has been extensively studied at the cellular level through electrophysiology and two-photon (2P) imaging, the response of non-neuronal cells, particularly microglia, as well as the effects of ICMS on blood-brain barrier (BBB) integrity remain poorly understood. In this study, we applied ICMS under 2P imaging in dual-reporter mice, with green fluorescent protein labeling microglia and a red fluorescent Ca^2+^ indicator labeling neurons. We also monitored vascular dye leakage to assess BBB integrity throughout the experiment. Using clinically relevant waveform parameters, we tested a range of current amplitudes. Surprisingly, we found that microglia responded rapidly, within 15 minutes of stimulation, by converging their processes (MPC) on areas of high neural activity. The prevalence of MPC increased with higher current amplitudes, but intriguingly, it did not correlate with the strength of the local electric field. Additionally, vascular dye penetration into brain tissue was higher in stimulated animals than in controls and increased with current amplitude. This study reveals a rapid microglia and BBB response to ICMS that has not been reported previously, underscoring the need for further research to fully characterize the biological response to ICMS and establish improved safety standards.

## Introduction

Motor and sensory deficits caused by peripheral or spinal nerve injury or neurodegenerative diseases are features of many neurological conditions. Brain-implanted microelectrode arrays (MEAs) have been used in animals and humans to record neural activity related to motor planning and execution. These signals can then be translated by a brain-computer interface (BCI) to control a robotic prosthesis ^1 2 3^. More recently, electrical intracortical microstimulation (ICMS) delivered via implanted MEAs has been used to elicit tactile sensation in patients by driving action potentials in neurons of the somatosensory cortex, providing haptic feedback to users of BCI-controlled prostheses ^4^. The principle of ICMS has been applied in other domains including hearing and vision and holds much promise for sensory restoration ^5 6 7 8^. Although studies have begun to illuminate the relationship between the strength of the neural response and ICMS parameters, the response of non-neuronal cells and in particular microglia – the resident immune cells of the brain – and effects on blood-brain barrier (BBB) integrity are still relatively poorly understood ^9^.

A key objective for studies of the cellular response to ICMS is establishing limits for safe and effective stimulation, which is of critical importance for improving BCI functionality. Our current understanding of the cellular response to ICMS is largely based on histological studies of stimulated tissue, often focusing on neurons ^10 11 12 13 14^. Early studies were carried out in the feline cortex using surface electrodes, and the categorization of the stimulation parameters as damaging or non-damaging was based on histological evidence ^11 12^. This empirical evidence was later used to formulate an equation for assessing the safety of given stimulation parameters, commonly known as the Shannon equation ^15^. This was an important step in the study of neural stimulation, but there are still many gaps in our understanding. The equation is based on data from surface electrodes and does not translate well to stimulation delivered via intracortical microelectrodes ^16 17^. It also only accounts for charge density and charge per phase, leaving out such stimulus parameters as: pulse rate, pulse duration, duty cycle, and total stimulation duration. Subsequent models with access to more data sets and utilizing machine learning approaches have attempted to broaden the applicability of such models but still only achieved an accuracy of 88.3% when compared to ground truth data ^18^. Critically, these models of tissue damage thresholds were based on histology, which provides only a discrete endpoint and little information regarding how the tissue response evolves over time.

The effects of alternating-current stimulation on microglia have been studied in modalities other than ICMS, including deep brain stimulation (DBS) and spinal cord stimulation. Histological studies have shown a reduction in inflammatory activation and an increase in the number of ramified microglia around stimulated DBS implants compared to unstimulated controls in normal rats ^19^ as well as in Parkinson’s disease model rats ^20^. Spinal cord stimulation for treating chronic pain also resulted in reduced microglia activation compared to unstimulated controls ^21 22^. These results are interesting in that they suggest electrical stimulation can modulate microglia activation. However, the mechanisms governing this are not understood. Previous work has been done in vitro to study the direct effect of electric fields on microglia. These studies have found that large direct current fields can orient microglia, direct their migration, and affect morphology, surveillance and gene expression ^23 24 25 26 27^. Previous modeling work has shown that the strength of the electric field drops off rapidly when using electrodes and stimulus parameters analogous to those used here, following an inverse square law ^28^. Therefore, any direct effects of stimulation should occur only in very close proximity to the stimulating electrode. The extent to which these findings apply to ICMS is also unclear, as there are large differences in stimulation locations, electrode sizes and stimulation frequencies. Two-photon (2P) microscopy has been used to characterize the real-time acute response of microglia to MEA implantation in the absence of ICMS ^29^ and found that the microglia respond rapidly to probe implantation on the timescale of minutes to hours. The response of microglia to ICMS on these timescales has not been examined.

ICMS may have effects on glial physiology beyond long-term changes in gliosis, such as altering neuron-microglia interactions. This is plausible given that it is known that microglia interact with neurons to prune synapses ^30 31^ and also play a critical role in neural development ^32 33 34 35^. The neural response to ICMS has been comparatively well studied as the sensations generated by ICMS are driven by neurons firing action potentials in response to electrical stimulation. The neural responses have been studied behaviorally ^36^ and electrophysiologically ^37^ as well as via 2P microscopy ^28 38 39 40 41 42^. These studies have established thresholds for the behavioral report and neural activation and have begun to define input-to-response relationships in terms of current amplitude, waveform shape and stimulation frequency.

The BBB response to ICMS is of primary interest in the present study as it has also been understudied compared to the neural response ^9^. MEA insertion is known to cause BBB disruption ^43^ from damage to the meninges and small vessels. This includes increased levels of blood proteins such as albumin and IgG around implanted MEAs ^44 45 46^. Vessel remodeling has also been observed around chronically implanted MEAs ^47^. Blood and tissue oxygenation as well as vessel dilation in response to ICMS have been studied ^48 49^, however there is little data on the direct effect of ICMS on BBB permeability. There is some evidence to suggest that electrical stimulation can increase BBB permeability based on early observations made using surface electrodes ^50 10^, however these few observations warrant much greater study given the rise in the clinical application of ICMS.

In the present study, we took advantage of transgenic mouse lines combining cx3cr1-GFP and thy1-RGECO mice via breeding to express GFP in microglia and a red Ca^2+^ indicator in neurons. To study the effects of ICMS current amplitude on BBB integrity, 3kDa-dextran Texas Red vascular dye was injected intraperitoneally after MEA implantation. The amplitudes of stimulation current tested were from 0 to 140μA for a total of 11 different current amplitudes. Tracking microglia on the timescale of minutes to hours during ICMS allowed us to observe the rapid and coordinated microglia process convergence (MPC) we report here. MPC is distinct from normal microglia process extension in that it involves the simultaneous convergence of processes from multiple microglia onto a central point ^30^. In the experiments described here, it occurred rapidly within 15 minutes following stimulation and was more prevalent at higher currents.

## Methods

### Animals

A total of ten mice (cx3cr1-GFP:thy1-jRGECO1a) originally obtained from the Jackson Laboratories and bred for this study (stock# 005582 and 030527). Mice 8 to 16 weeks old were enrolled in this study. All experiments were performed in accordance with the institutional guidelines set by the Institutional Animal Care and Use Committee of the University of Pittsburgh.

### Electrode array preparation

16-channel 5mm single shank NeuroNexus MEAs (A1×16-5mm-50-703-CM16LP) with iridium sites were used for this study. Before implantation, the iridium electrode sites were electrochemically activated to iridium oxide to increase the charge injection limit by the application of 300 cycles of a –0.85V to 0.75V, 0.5Hz to 1Hz square wave potential based on the manufacturer protocol. Probes were subsequently characterized via cyclic voltammetry, electrochemical impedance spectroscopy and voltage transient measurement of single pulses following the stimulation waveform to verify the oxidation state of the Ir sites as described previously ^28^.

### Surgery and electrode array implantation

Surgical preparations were as previously reported ^29^. Briefly, mice were anesthetized with 75mg/kg ketamine and 10mg/kg xylazine intraperitoneally throughout the surgical procedure and subsequent imaging. The depth of anesthesia was monitored through respiration rate and toe pinch response. A craniotomy was performed without durotomy. The 5mm 16-channel single shank MEA was implanted into the cortex under 2P at 30° from horizontal using a microdrive (MO-81, Narishige, Japan) to a depth of approximately 200μm. 100μL of 10mg/mL 3kDa-dextran conjugated Texas Red dye was administered intraperitoneally following probe implantation. For ATP studies, saline supplemented with10mM ATP was used to fill the space between the 2P objective and craniotomy beginning ∼1hr prior to array implantation and throughout the duration of the experiment.

### Stimulation paradigm

Electrical stimulation of currents ranging from 5μA – 140μA in amplitude were delivered to a single electrode site in increasing order in the mouse cortex via a NeuroNexus MEA connected to an Autolab potentiostat. For each condition, the 200μs cathodic leading, 100μs inter-pulse interval and 400μs anodic trailing charge-balanced waveform was administered at 50 Hz for 6 repetitions of 1s on 5s off (16.7% duty cycle). These conditions were held constant and only the current level was varied (Fig. 1A). The waveform and stimulation parameters tested in this study were chosen to align with clinically relevant ICMS used to evoke sensory percepts in somatosensory ^4 51^ and visual ^8 5^ cortex. Four mice were in the sequential current group, receiving a sequence of increasing current stimulation: 0, 5, 10, 20, 30, 40, 60, 80, 100, 120, 140μA. Two mice received stimulation with a random order of current amplitude. Currents 5μA – 40μA were binned into ‘low currents’ and 60μA – 140μA were binned into ‘high currents’. Low currents were administered in a random order followed by high currents administered in a random order, without replacement, such that each stimulus condition was delivered once per animal. Mice that received randomized current administration were grouped with all other mice for analysis as no significant differences in microglial or neuronal activity were observed. Two mice were in the ATP group receiving sequential stimulation in the presence of ATP and two mice were in the no-stimulation control group receiving probe implantation and imaging following the same time course but without stimulation (Fig. 1C). This no-stimulation control group was used to distinguish the effects of ICMS as distinct from those of MEA implantation.

**Figure 1.**
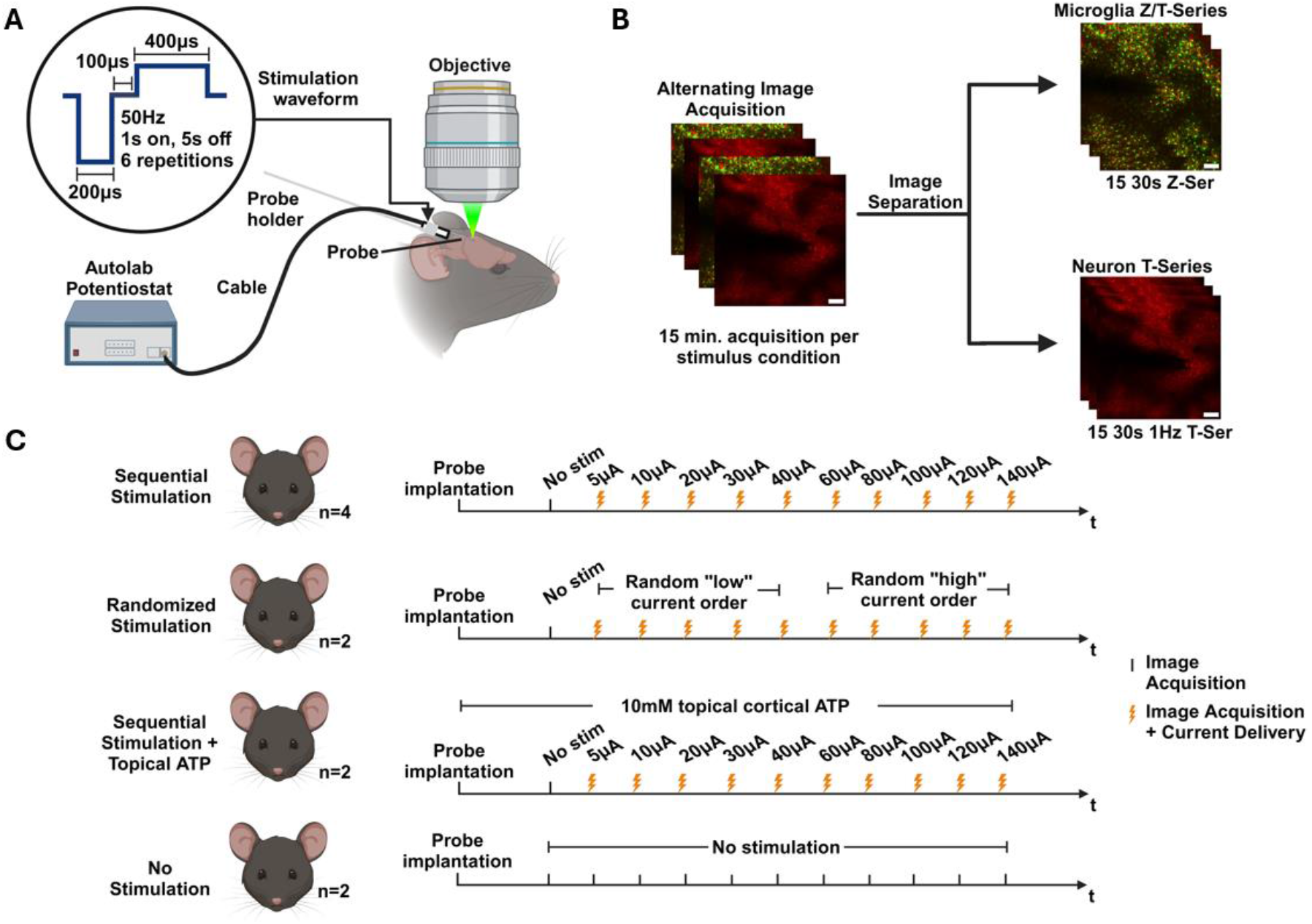
Experimental Paradigm for Assessing Neuron and Microglia Response to Microstimulation. **A)** Dual labeled cx3cr1-GFP:thy1-RGECO mice expressing GFP in microglia and red calcium indicator in neurons were anesthetized and acutely implanted with an IrOx coated MEA. Microstimulation pulses were delivered to a single electrode via an Autolab potentiostat at a current range of 0 to 140μA with a 200μs cathodic, 100μs interstimulus, 400μs anodic charge balanced waveform delivered at 50 Hz on a 1 sec. on/5 sec. off duty cycle for 6 repetitions. **B)** 2P imaging began at the initiation of each stimulation condition alternating between 30 seconds of a 1040nm 1 Hz T-series to image neuronal activity and 30 seconds of 920nm 30μm thick 3μm step size Z-series to image microglia morphology. After data collection, images were separated into neuronal and microglial sequences. For neurons, 30 seconds of 1 Hz Ca^2+^ imaging per 1 min. for 15 min. during and following stimulation was obtained. For microglia, one 30μm Z stack per minute for 15 minutes following stimulation was obtained and collapsed to generate a 15 image Z/T-series at 1 frame per minute. Scale bars are 100μm. **C)** Data from 4 experimental groups was collected. The sequential stimulation group consisted of 0,5,10,20,30,40,60,80,100,120,140μA stimulation in that order, n=4 mice. The random stimulation group consisted of 0μA followed by the 5-40μA conditions in random order followed by the 60-140μA conditions in random order, n=2 mice. The topical ATP condition was identical to the sequential condition but for the addition of 10mM ATP to the saline used to hydrate the brain. The no stimulation condition consisted of repeated 0μA stimulation following the same time course as other experimental groups, n=2 mice.

### 2P imaging and image processing

Fluorescence images were acquired with a 2P microscopy system (2P Plus, Bruker Nano, Inc.) equipped with a 16x 0.8 NA objective lens (Nikon, Inc.) and tunable laser (Insight X3, Newport Spectra-Physics, Inc.). All images were acquired at 512×512 pixel resolution, 1.3x zoom, and 2.4μs dwell time, resulting in a field of view (FOV) of 867.46μm x 867.46μm and a frame period of 0.987s. The FOV was roughly centered on the stimulating electrode in x, y and z planes, with the z depth approximately 150μm below the pia mater. Microglia and vasculature were imaged using 920nm wavelength excitation and neuronal images were acquired using 1040nm excitation. For each stimulation, imaging began at stimulus onset and continued for 15 minutes. For microglial and vascular images, a 30μm Z-stack (Z-series) with 10 slices (3 μm spacing) was acquired once per minute for 15 minutes pre-, peri- and post-stimulation (30s pre-stimulation). Neuronal times series (T-series) were acquired for 30s at 1 Hz frequency in a single plane interleaved with the Z-stack imaging of microglia every minute for 15 minutes peri- and post-stimulation. (Fig.1B). For the no-stimulation control animals, the same imaging parameters were used and followed the same schedule as stimulated animals, but in all conditions 0μA was delivered.

### Image Processing

Images were pre-processed in ImageJ to separate Z- and T-series imaging for microglia and neurons, respectively. All images were then imported into MATLAB where motion correction and intensity correction were performed using custom scripts. MPC regions of interest (ROIs) were defined as the point where processes from two or more microglia converged, including the area extending out from this point to the soma of the converging microglia. Example MPC events can be seen in Supplemental Videos S2,3,5,6 and in Fig. 3A. Example MPC ROIs can be seen in Fig. 4C and Fig. 5B. were applied to eliminate portions of vasculature that remained. Following vessel mask subtraction, neuronal T-series average intensity and standard deviation projections were used to quantify neuropil activation.

**Figure 2.**
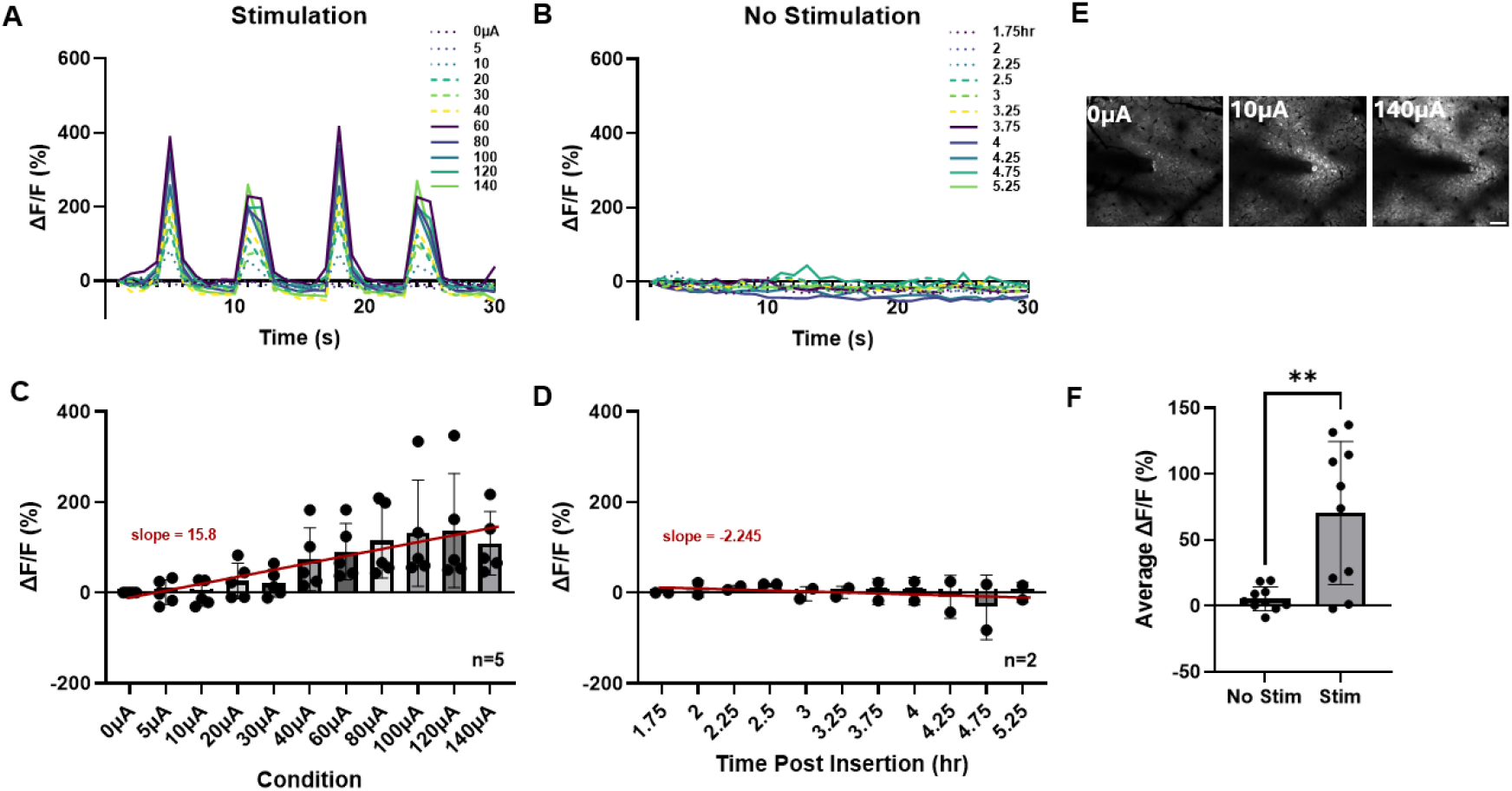
Neuronal response to increasing microstimulation current amplitude. **A)** Average neuronal RGECO fluorescence intensity increase (%dF/F) in the first 30 sec. following stimulation onset. n=1 mouse. **B)** Average neuronal RGECO fluorescence intensity increase (%dF/F) in the first 30 sec. in the no stimulation control group. n=1 mouse. **C)** Average %dF/F for each stimulation condition. P<0.0001 slope significantly non-zero by linear regression. n=5 mice. **D)** Average %dF/F for each timepoint in the no stimulation control group. P>0.05 by linear regression. n=2 mice. **E)** Average projection of the first 30 sec. following stimulation onset for example 0, 10 and 140μA current amplitudes. Scale bar is 100μm. **F)** Average %dF/F for each condition in the stimulation and no stimulation group. P<0.0001, unpaired t-test, n=10 conditions.

**Figure 3.**
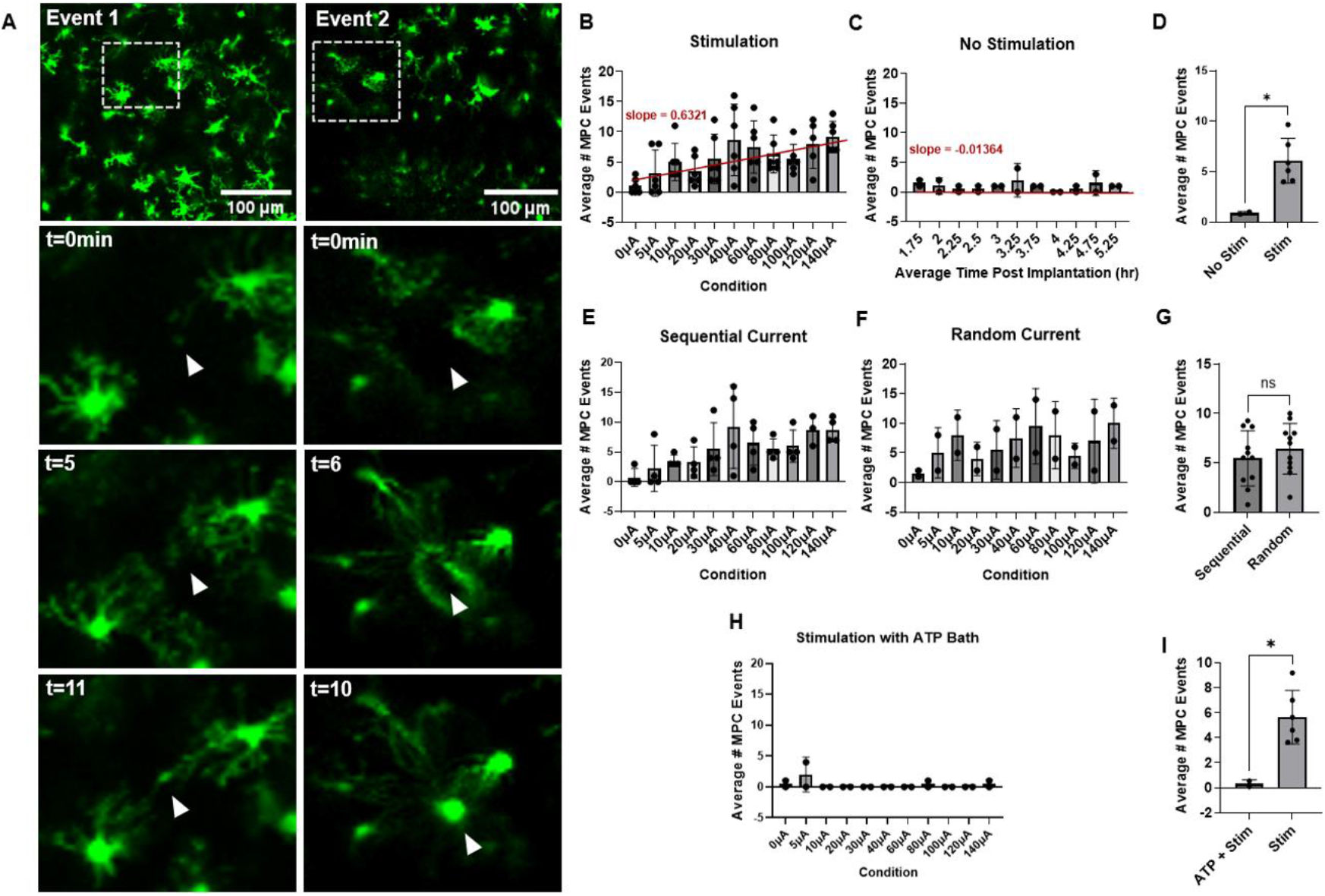
Microglial response to increasing microstimulation current amplitude. **A)** Example microglia process convergence events. Left and right columns represent two separate convergence events. The top panel is a zoomed-out view showing the convergence ROI displayed below (dashed boxes). White triangle indicates approximate point of convergence. **B)** Average number of convergence events in the 15 minutes following stimulation onset for each stimulation condition. n=6 mice. **C)** Average number of convergence events in the 15 minutes following each timepoint in the no stimulation control group. n=2 mice. **D)** Average number of convergence events per condition for each mouse between the no stimulation control group and stimulation condition. P<0.05 t-test. n=2,6 mice. **E)** Average number of convergence events in the 15 minutes following stimulation onset for each stimulation condition in the sequential current group. **F)** Average number of convergence events in the 15 minutes following stimulation onset for each stimulation condition in the random current group. **G)** Average number of convergence events per condition for each mouse between the sequential current and random current groups. **H)** Average number of convergence events in the 15 minutes following stimulation onset for each stimulation condition in the stimulation with ATP bath group. **I)** Average number of convergence events per condition for each mouse between the stimulation with ATP bath and stimulation groups. P<0.05 unpaired t-test n=2,6 mice.

**Figure 4.**
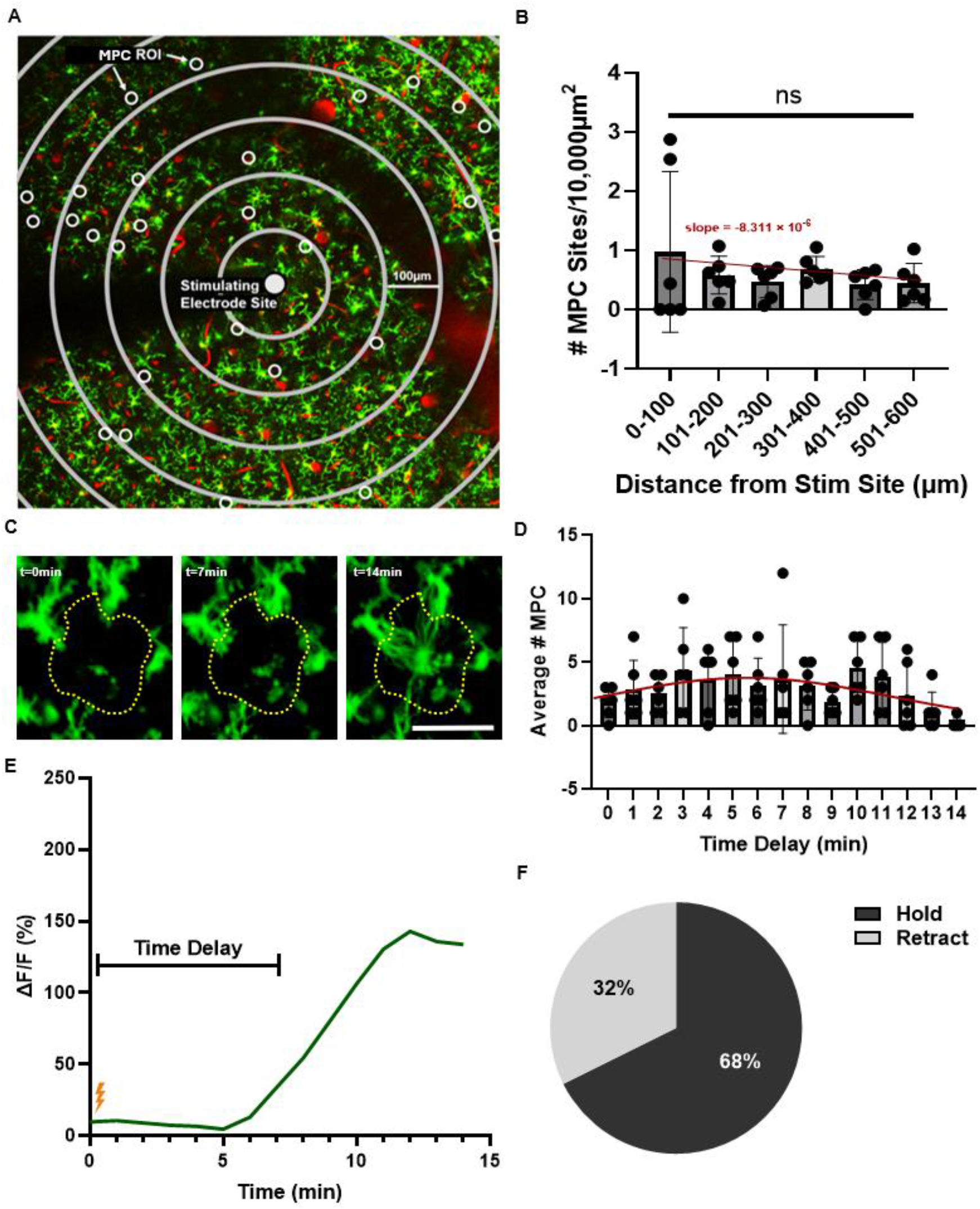
Distribution and time course of microglia convergence events. **A)** Example field of view showing locations of microglia convergence events in grey, the stimulating electrode site in white and the concentric rings used for the analysis in B spaced 100μm apart. **B)** Average number of convergence sites per 10,000μm^2^ in each 100μm bin, P>0.05 1-way ANOVA. **C)** Example time course of a microglia convergence event. Example images are 0, 7 and 14 minutes post stimulus. The %dF/F within the yellow dotted line is plotted below. Scale bar is 50μm **D)** Average number of convergence event onset times in the 15 minutes following stimulation onset. **E)** The %dF/F within the ROI denoted by the yellow dotted line shown in C. **F)** Proportion of MPC which persisted for the duration of the experiment (dark grey) vs retracted at some point during the experiment (light grey). n=6 mice.

**Figure 5.**
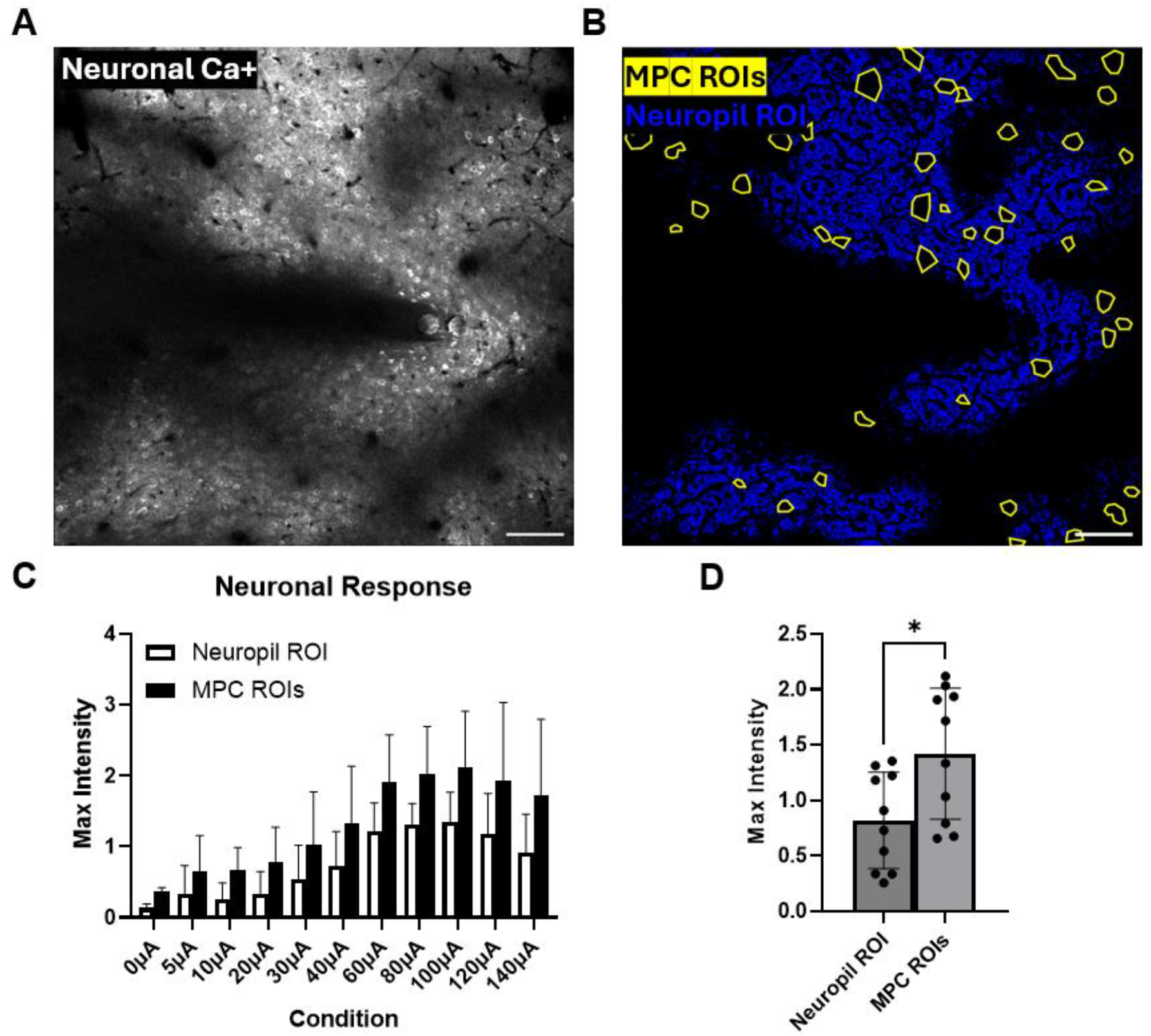
Microglial-neuron interaction during microstimulation. **A)** Standard deviation projection of the RGECO signal in the 30 sec. following stimulation across the whole field of view, showing the areas of neuropil activation as well as areas of poor imageability. **B)** Masks applied to the image in A showing convergence ROIs in yellow and the imageable neuropil in blue. Images shown in panels A and B both have vasculature masked out. Scale bars in A and B are 100μm. **C)** Average maximum RGECO fluorescence intensity within the convergence ROIs and within the neuropil ROIs for each stimulation condition. **D)** Average maximum RGECO fluorescence intensity in all stimulation conditions averaged across mice between the convergence ROIs and the neuropil ROIs. P<0.05 t-test.

### ROI-based neuropil analysis

To assess the link between neuronal activity and MPC events, neuropil intensity was measured within MPC ROIs. Maximum neuropil intensity in each ROI was measured in MATLAB-processed images without vascular artifacts or shadows and maximums were averaged across all ROIs per animal. The same approach was implemented for the entire imaging field excluding MPC ROIs, vasculature, and un-imageable regions (probe shadows, etc.) for each animal.

### Distance analysis

Concentric rings every 100μm were applied to images and centered at the stimulating electrode site for distance analysis. The number of MPC ROIs per annulus were counted and normalized to the area of the annulus. In addition, average neuropil intensity was measured per annulus.

### Statistical analysis

Linear regressions shown are simple linear regression without fixed intercepts, significance is slope different from 0 with alpha equal to 0.05. For determining the response threshold for neurons and microglia, one-way ANOVA followed by post-hoc t-tests with alpha equal to 0.05 were used. The trendline for MPC in each 1 min. time bin following stimulation was a Gaussian fit to the histogram. Unpaired t-tests were used for significance comparisons of aggregated data, with an alpha of 0.05.

## Results

### Increasing microstimulation current leads to increasing neural response

We quantified the neuronal response to a range of microstimulation currents to determine the timing, strength and current response relationship. Stimulation induced a rapid increase in neuronal Ca^2+^ in response to each 1s burst of stimulation, which decayed to baseline within 2 seconds during the 5s interval between bursts (Fig. 2A). In the no-stimulation control mice, the Ca^2+^ signal remained stable in the first 30 seconds of imaging and this trend continued for the duration of the experiment (Fig. 2B). The average %dF/F increased with increasing current amplitude (Fig. 2C) whereas in the no-stimulation control mice the average %dF/F was small and stable for the duration of the experiment (Fig. 2D). A linear regression of %dF/F on current amplitude was significantly different from 0 (p<0.0001) with a slope of 15.8 and R^2^ of 0.885, whereas for the no-stimulation condition it was not significantly different from 0, the slope was −2.245 and the R^2^ was 0.3293. Example images in Figure 2E show the increasing Ca^2+^ reporter fluorescence with increasing current amplitude. The difference in %dF/F between stimulated and no stimulation control mice was significant across conditions (Fig. 2F, p<0.0001). This level of graded response is similar to what has been observed in other 2P studies using similar ICMS parameters ^52 53^. We performed a one-way ANOVA on the data in Fig. 2C and found that there was a significant effect of current amplitude (p<0.01). Further post-hoc tests revealed that all conditions 40μA and above were significantly different from the 0μA condition.

### Microglia respond to microstimulation by rapid and coordinated process convergence

To dissociate the acute microglial response to probe implantation from the microglia response to electrical stimulation, we waited at least 90 minutes post-implantation before stimulating, monitoring the acute microglia response to probe implantation until it had subsided. To further remove any influence of implantation on our results, we excluded the microglia directly interfacing with the probe shank from our analyses. Following the application of ICMS, we observed multiple occurrences of MPC, the rapid and coordinated extension of processes from multiple microglia converging towards a central point (Fig. 3A, Supplemental Movie S1). These events were observed following initial stimulation and increased in prevalence as current amplitude increased (Fig, 3B), whereas they were quite rare in the no stimulation control animals and remained at a stable low level for the duration of the experiment (Fig. 3C). Comparing the average number of MPC events per condition between the stimulation and no stimulation groups, we found significantly more MPCs in the stimulation group (Fig 3D, p<0.05). In addition to the sequential current group (Fig. 3E), we also tested a random current application group (Fig. 3F) and found that they were not significantly different from each other (Fig. 3G). We therefore combined these groups for the graph shown in figure 3B and for subsequent analyses.

### MPC is current dependent

To assess whether there was a relationship between current amplitude and the prevalence of MPC, similar to the relationship between %dF/F and current amplitude for neurons, we fit a linear regression to the number of MPC in the stimulated condition and found that it was significantly different from 0 (p<0.0001) with a slope of 0.6321 and R^2^ of 0.2415, whereas for the no stimulation condition it was not significantly different from 0, the slope was −0.01364 and the R^2^ was 0.001782. Due to the relatively low R^2^ for the linear fit in the stimulated condition and the apparent plateau in the number of MPC at 40μA, we also performed an exponential plateau nonlinear fit with a corresponding R^2^ of 0.7563. The slope of the linear regression in the sequential current group (Fig. 3E) was 0.6364 and the R^2^ was 0.2516. In the random current group (Fig. 3F) the slope was 0.4727 and the R^2^ was 0.1331. We performed a one-way ANOVA on the data in Fig. 3B and found that there was a significant effect of current amplitude. Further post-hoc tests revealed that 10μA and above were significantly different from the 0μA condition (p<0.05).

### MPC is dependent on purinergic signaling

We performed an additional set of experiments where we abolished local purine gradients by adding ATP (10 mM) to the saline bath applied to the brain, similar to the methods described previously ^30 54^. Under these conditions, MPC was almost entirely absent for the duration of the experiment across all current conditions (Fig. 3H). This was significantly different (p<0.05) from the number of MPC events observed without the application of ATP (Fig. 3I).

### MPC is not a direct electric field effect

We assessed the prevalence of MPC relative to the distance from the stimulating electrode. Locations of MPC across a whole FOV relative to the stimulating electrode are shown, demonstrating that MPC occur with relatively equal prevalence across the FOV and are not restricted to near the stimulating electrode (Fig. 4A). We binned the field of view into 100μm concentric rings centered on the stimulating electrode and calculated the number of MPC as a function of imageable area in each ring (Fig. 4B). We found that there was not a significant relationship with distance from the electrode and that the slope of the best fit line was small and not significantly different from zero, at least within the field-of-view.

### MPC occurs rapidly following stimulation and is dynamically reversible

We quantified the timing of MPC by calculating the time in minutes to MPC initiation within each ROI. Example images used to generate a fluorescence trace to determine the time of MPC initiation are shown in Figure 4C. The time of initiation for all MPC are plotted in Figure 4D. Most MPC began within 15 minutes following stimulation. An example trace is shown in Figure 4E. The mean time to initiation for all MPC was 6.39 minutes with a standard deviation of 3.67 minutes. This timing profile is on the order of process extension and contact with the probe observed directly following implantation ^29^. We also observed that 32% of MPC retracted at some point during the experiment and 68% remained in a converged state (Fig. 4F).

### MPC occurs predominantly at sites of elevated neural activity

We compared the strength of the neural response to ICMS at MPC sites to the strength of the neural response to ICMS across the rest of the imageable FOV. Figure 5A shows an example of the neural response to stimulation and Figure 5B shows the two masks used to determine the relative neural response within MPC regions (yellow) and in the rest of the imageable neuropil (blue). We found that the maximum %dF/F was consistently higher in MPC ROIs than in the rest of the observable neuropil (Fig. 5C) and that this difference was significant across conditions (Fig. 5D).

### Intracortical microstimulation increases blood brain barrier permeability

By injecting a 3kDa dextran conjugated vascular dye at the beginning of each experiment, it was possible to measure the degree of BBB permeability throughout the experiment. To track the level of tissue penetration from the bloodstream of the fluorescent 3kDa dextran over time, we masked out the vasculature present in the FOV, leaving only parenchyma, and assessed the normalized mean intensity of the vascular dye present in the tissue. To control for vascular damage due to probe insertion, only tissue 500μm and further from the probe was quantified. Texas Red intensity in the tissue across time in the stimulation and no stimulation groups is shown (Fig. 6A). To determine the degree of change, we fit a line to the data from each group (Fig. 6B) and found that the slope for the no stimulation group was small and not significantly different from zero whereas for the stimulation group the slope was larger and significantly positive (p < 0.05).

**Figure 6.**
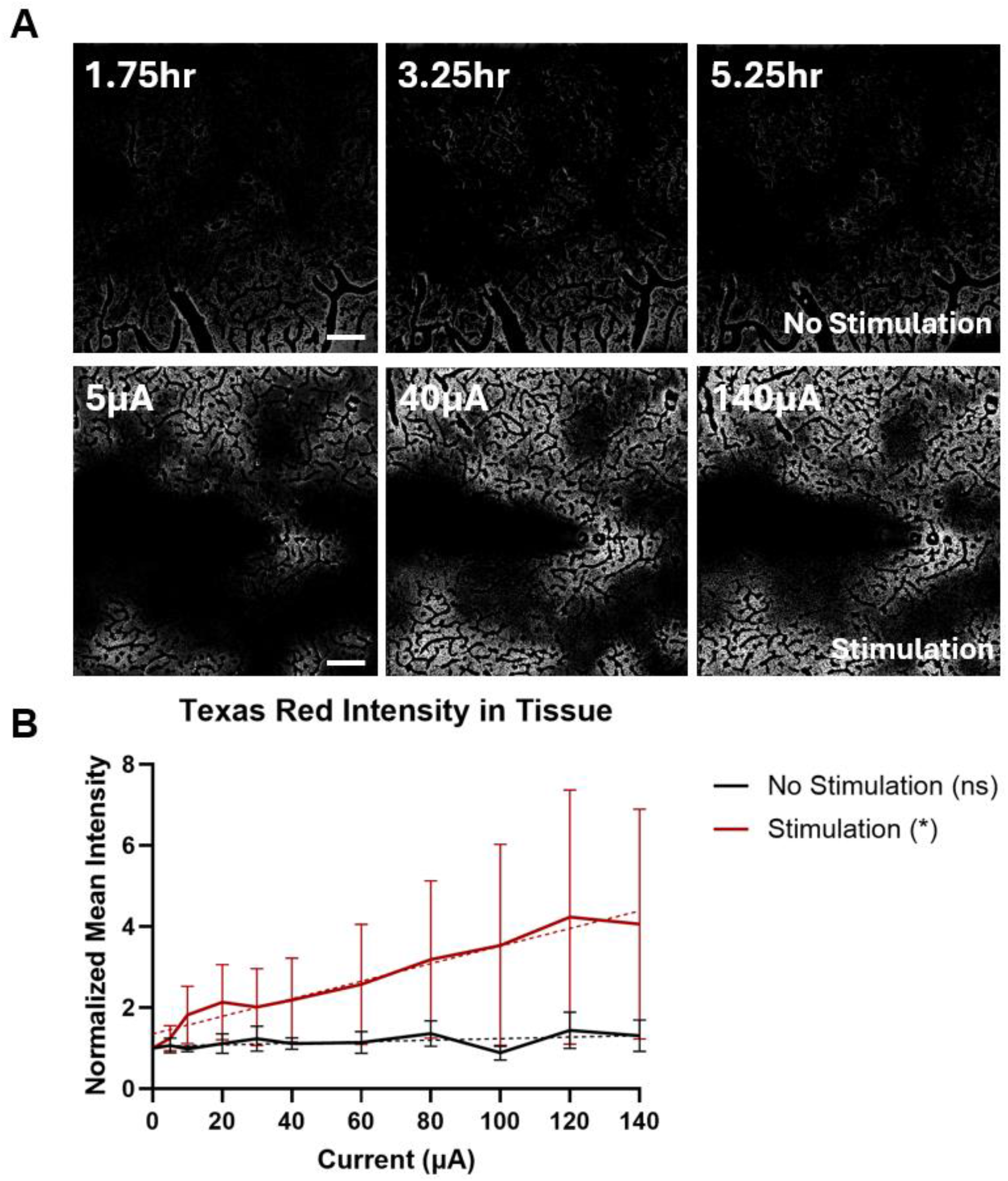
Blood brain barrier permeability response to increasing microstimulation current amplitude. **A)** Example time course of 3kD Dextran-Texas red leakage into the tissue in the no stimulation condition (top row) and the stimulation condition (bottom row) time post implant in the no stimulation condition corresponds to the time the stimulus was administered in the stimulation condition. A mask was applied to remove vessel fluorescence from the images. Scale bars are 100μm. **B)** Average Texas red fluorescence intensity in the tissue 500μm and further from the probe (excluding the vessel mask) for each condition in the stimulation (red) and no stimulation (black) conditions. P<0.05 slope greater than 0 in the stimulation condition, slope not significantly different from 0 in the no stimulation condition. Linear regression n=5,2 mice.

## Discussion

ICMS can be a powerful tool for both basic neuroscience as well as clinically relevant applications. Although its primary purpose is to drive a neural response, it can have off-target effects on non-neuronal cells and vasculature. It is critical to understand the effects of ICMS on non-neuronal cells and the BBB in order to better apply it effectively and safely. Here, we used 2P to elucidate the rapid, current dependent microglia and BBB responses to ICMS across a range of current amplitudes. We show MPC wherein multiple microglia converge their processes on a central point of high neural activity within 15 minutes of stimulation and that these events are more prevalent at higher current amplitudes. We also show that BBB permeability is elevated following stimulation, and that permeability increases as current amplitude increases. We found that MPC occurred across the FOV and was not restricted to near the stimulating electrode, and furthermore, there was no relationship between distance from the electrode and the prevalence of MPC. Given the lack of correspondence between MPC and proximity to the stimulating electrode, we sought an explanation for the MPC trigger other than the strength of the local electric field around the microglia. Both neurons and microglia across the FOV responded to stimulation, and both the neural response as well as the number of MPC increased as the stimulation current increased. We hypothesized that the trigger initiating MPC was mediated not by the strength of the direct electric field but by the strength of the local neural response. We found evidence supporting this hypothesis in that MPC occurred with roughly equal prevalence close to the electrode and distally, where the electric field generated by stimulation was negligible. Furthermore, we show that MPC preferentially targets areas where the neuronal response is particularly high. This suggests that MPC is not dependent on the strength of the local electric field but on a secondary effect, likely the activity of neurons driven by the stimulus which, due to synaptic transmission, is much more far-reaching than the electric field itself ^37 42^.

The present study highlights the far-reaching effects of ICMS on diverse cell types and physiology and the rapidity with which it can affect non-neuronal cells. At present, our data suggests that keeping the stimulation current amplitude below 10μA under these conditions should reduce ICMS induced MPC and it is possible this may be the best approach to maximize long term MEA performance. Keeping the number of rounds of stimulation as low as possible to achieve a therapeutic effect would also reduce MPC. Conversely, increasing current amplitude and number of stimulation rounds would increase MPC. This may even be therapeutically advantageous under certain circumstances, as there is some evidence to suggest that electrical stimulation can decrease gliosis and potentially increase device performance ^55 56 57^, although the exact mechanisms remain poorly understood and the impact of ICMS on long term gliosis and device performance and longevity have yet to be fully determined. It is possible that ICMS induced MPC may not be a product or a cause of increased gliosis per se, but rather a more nuanced response that remodels the neural network most strongly activated by ICMS. Keeping both current amplitude and number of stimulation rounds as low as possible should also keep the ICMS induced BBB leakage to a minimum, and this may be the best approach to ensure the health of the tissue. It is also possible that ICMS could be used therapeutically to open the BBB locally around electrodes to allow circulating therapeutic agents into the brain tissue. Future chronic studies focused on determining the ultimate fate of neurons and microglia as well as vasculature involved in ICMS induced MPC should address these questions more thoroughly.

The fact that abolishing local ATP gradients by the application of ATP uniformly across the whole craniotomy also abolished MPC in our experiments lends weight to the hypothesis that release of ATP by hyperactive neurons was driving the MPC we observed. Mechanistic studies of microglia process extension and microglia-neuron interactions have implicated P2Y12, an adenosine binding G-protein coupled receptor highly expressed on microglia, as being critical for microglia process extension ^58 59^. The results from our ATP experiments were similar to experiments performed in slices when extracellular ATP was uniformly applied as well as when using tissue from P2Y12 knockout mice, as in all cases MPC was abolished ^60 30 54^. It has been shown that various stimuli such as reducing extracellular Ca^2+^, applying glutamate, kainic acid or repeated stimulation of single neurons via a patch clamp electrode all induce MPC ^30 61 62^. Different mediators and downstream interactions with neurons were observed for different triggers. Glutamate and kainic acid-induced MPC was NMDA, CX3CR1 and IL1-β dependent and targeted dendrites, whereas repeated patch clamp stimulation targeted axons. In all cases P2Y12 and ATP were implicated, as in the present study. These studies also suggest a potential neuroprotective effect of MPC whereby neurons directly contacted by microglia showed reduced depolarization following repeated stimulation ^62^. In another study, loss of CX3CR1 signaling resulting in loss of MPC led to a more intense seizure response to kainic acid treatment ^61^. The role of secondary mediators such as CX3CR1, IL1-β or others in ICMS-induced MPC has yet to be determined.

One of the striking things about the MPC we observed here was how rapidly it occurred following stimulation. It is also interesting to note that the stability of MPC was not uniform following formation, as some would persist for the duration of the experiment whereas others would retract. This suggests that local ATP gradients may not be stable over the time course of these experiments, or that there may be additional cues governing whether downstream processes such as synaptic pruning are engaged ^63^ or whether microglia return to their previous surveilling state ^64^. Taken together, these results begin to shed light on the non-neuronal responses to ICMS which occur rapidly and likely involve microglia-neuron interactions.

In addition to the neuron and microglia responses, we found that BBB permeability increased following repeated stimulation at increasing current amplitudes. This occurs rapidly following stimulation onset, on the time scale of tens of minutes to hours. Although the 3kDa dextran used here is smaller than many blood proteins, there are several peptides and small molecules of around 3kDa and smaller that are present in the bloodstream and can be detrimental if they penetrate into the brain parenchyma, including bradykinin, substance P and lactate ^65 66 67^. This suggests that repeated application of ICMS can increase the permeability of the BBB to molecules of at least 3kDa in size, of which many are present in the bloodstream. This increased BBB permeability could have many downstream consequences for the tissue and warrants further investigation in future studies. Previous studies of BBB damage have shown MPC onto damaged vessels and that convergence can facilitate BBB closure ^68^, which should be more thoroughly investigated in the context of ICMS.

Much future work needs to be done to elucidate the mechanistic nature and downstream consequences of the effects we observe here. In the present study, we imaged over a wide FOV which limited our spatiotemporal resolution such that we were not able to capture direct sub-cellular microglia-neuron interactions. Future studies with greater spatiotemporal resolution could capture subcellular interactions of microglia contacting neuronal synapses and somas and the ultimate fate of these neurons and microglia. The application of propidium iodide ^69^ or other membrane impermeable dyes could help determine whether MPC is occurring at damaged or dying neurons by seeing whether these neurons have particularly permeable cell membranes, which would suggest they may have been damaged by excitotoxicity resulting in increased membrane permeability. This could mediate the release of factors such as ATP that could serve as a signal for nearby microglia to begin MPC. Future studies employing ATP fluorescent sensor such as iATPSnFRs ^70^ could confirm these results and further elucidate the dynamics of ATP mediated MPC in the context of ICMS, as well as define the effects of ICMS on ATP dynamics generally.

## Supporting information

S2_m1_40uA_ROIA_microglia

S1_m1_40uA_ROIA_neurons

S6_m1_100uA_ROIC_merge

S5_m1_100uA_ROIC_microglia

S4_m1_100uA_ROIC_neurons

S3_m1_40uA_ROIA_merge

## Notes

### Competing Interest Statement

The authors have declared no competing interest.

### Summary of Updates

Introduction updated to discuss safety limits in more detail; Figures updated to include missing scale bars.

